# ProteoMeter: A pipeline for integrating multi-PTM and limited proteolysis data to reveal modification-structure coupling at the residue level

**DOI:** 10.1101/2025.07.14.664747

**Authors:** Jordan C. Rozum, Amy C. Sims, Xiaolu Li, Snigdha Sarkar, Tong Zhang, John T. Melchior, Danielle Ciesielski, David D. Pollock, H. Steven Wiley, Wei-Jun Qian, Song Feng

## Abstract

Systemic perturbations trigger extensive changes across the proteome–altering protein abundance, post-translational modifications (PTMs), conformational states, and complex assembly. Interpreting these effects demands computational pipelines capable of integrating diverse proteomics modalities, such as multi-PTM profiling, limited proteolysis mass spectrometry (LiP-MS), and cross-linking mass spectrometry (XL-MS), within a unified and interoperable framework. Because instrument data are quantified at the peptide level, mapping these measurements to individual residue or modification site is essential for biologically meaningful interpretation. We introduce `ProteoMeter`, an open-source Python library designed to integrate multi-modal proteomics datasets and map them to single-residue resolution using a standardized coordinate framework. We showcase its capabilities in a combined multi-PTM and LiP-MS analysis profiling the proteomic response to human coronavirus 229e (HCoV-229E) infection.`ProteoMeter`is actively maintained and is freely available–including all source code and figure-generation scripts–at the following repository: https://github.com/PNNL-Predictive-Phenomics/ProteoMeter.

## Introduction

Proteome-wide investigations have revealed that systemic perturbations–such as infection, stress, or treatment– can induce intricate changes in protein abundance, post-translational modifications (PTMs), conformational dynamics, and protein complex assembly. Capturing and interpreting these multi-faceted alterations demands advanced experimental techniques, including multi-PTM profiling, limited proteolysis mass spectrometry (LiP-MS), and cross-linking MS (XL-MS). While these modalities collectively offer a comprehensive view, they remain analytically siloed, with data quantified at the peptide rather than the residue level, introducing challenges in deriving biologically meaningful insights.

Multi-PTM proteomics and LiP-MS in particular have emerged as powerful proteomics tools, capable of mapping multi-type PTMs and conformational landscapes respectively across complex proteomes. In multi-PTM profiling, cysteine oxidation, phosphorylation, acetylation, and other PTMs are measured simultaneously from the same samples, which is valuable in unraveling the functional roles of different PTMs and their co-regulation [1]. In LiP-MS, half-tryptic peptides are generated upon limited protease digestion of solvent-exposed regions in folded proteins, with peptide-level quantification serving as a proxy for structural shifts [2, 3].

To date, computational tools for PTM [4, 5] and LiP-MS [6, 7, 8] data remain largely modality-specific and lack integrated support for PTM-informed structural inference. Those that enable *ad hoc* combining of analyzed LiP-MS results and published PTM information generally do not offer methods for quantifying changes in enzymatic cleavage patterns at the site level, instead emphasizing changes in peptide distribution [9]. There is also a scarcity of proteomics data analysis pipelines implemented as a Python library, which impedes interfacing with Python’s large and rapidly evolving proteomics machine learning ecosystem.

`ProteoMeter` bridges these gaps by offering an open-source Python toolkit that integrates multi-modal proteomics at the protein, peptide, and residue levels; aligns peptide-level quantifications to specific residues or modification sites; and supports downstream statistical, structural, and functional analyses. By standardizing configuration, analysis, and outputs across data types, `ProteoMeter` enables improved interoperability and biological interpretation. In this application note, we highlight the key features of our tool and demonstrate its utility in a multi-PTM/LiP study of human lung fibroblasts (MRC5) infected with human coronavirus 229e (HCoV-229e).

## Results

### A new tool for residue-level integrative proteomics

The`ProteoMeter` workflow is illustrated in Figure 1. In brief,`ProteoMeter` requires peptide-level and protein-level intensity quantification (PTM-annotated peptides or trypsin-only and LiP peptides) in tabular format, as well as user-provided metadata as inputs. It outputs protein-level, peptide-level, and residue-level quantification that is normalized between samples (using median centering or sample-matched normalization), batch-corrected, and (optionally) protein-abundance-corrected.`ProteoMeter` automatically computes pair-wise t-tests, multi-way ANOVA, and false discovery control according to a configurable parameters file. Advanced users can execute individual parts of the workflow with further customization options.`ProteoMeter` is freely available to install in Python via PyPI https://pypi.org/project/proteometer. The source code is posted on GitHub at https://github.com/PNNL-Predictive-Phenomics/ProteoMeter. We include the processing scripts used to generate the results of this manuscript as a demonstration in the `ProteoMeter` source repository.

**Fig. 1.**
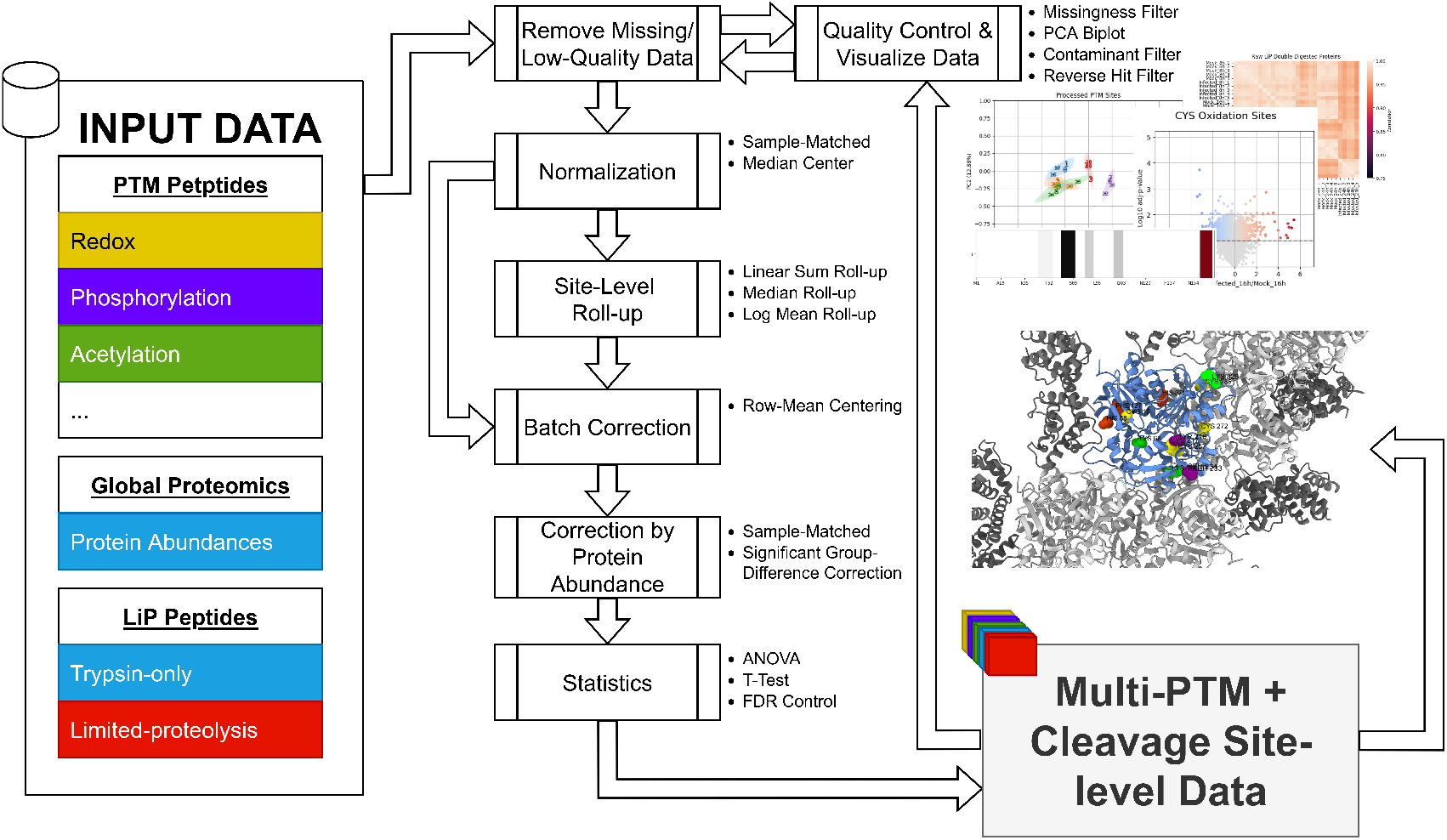
The `ProteoMeter` pipeline for PTM and LiP data. The program takes as input peptide-level quantification data from PTM and LiP datasets, as well as global protein abundances. The user may specify samples to exclude based on their assessment of quality control plots generated by `ProteoMeter`. Selected samples are normalized and rolled up to the site level. Next, batch and abundance corrections are applied, and `ProteoMeter` performs statistical evaluations as defined by the user in the configuration file. Processed protein-level, peptide-level, and site-level data frames are returned to the user that can be saved in tabular (e.g. comma-separated value) format or used to visualize proteins in external programs. `ProteoMeter` provides functions for visualizing outputs, such as volcano plots and barcode plots.

**Fig. 2.**
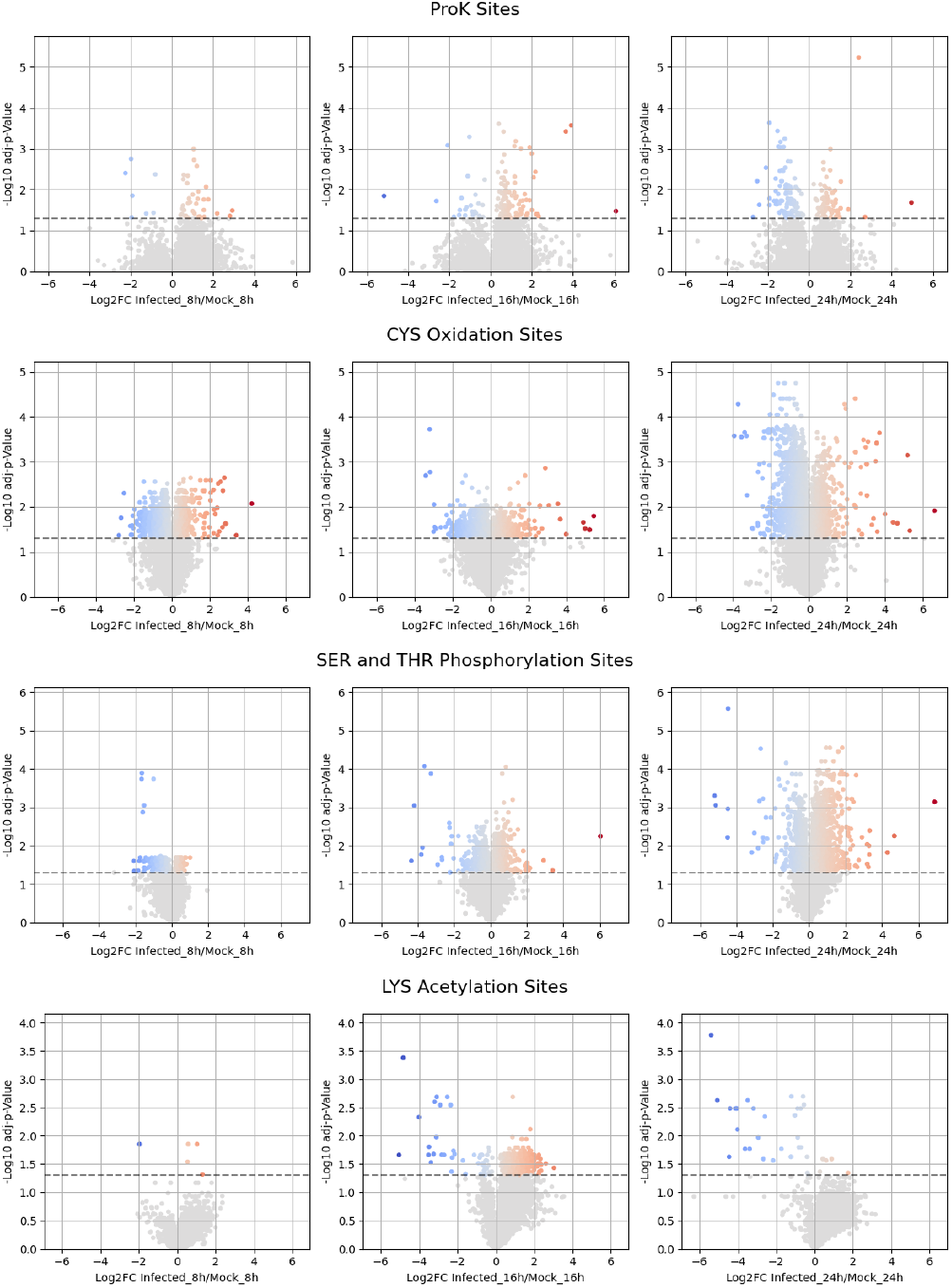
Volcano plots for site-level quantification. Points are colored on a blue-red gradient if above the adjusted p-value cutoff of 0.05. Each row represents a different site-type, as labeled. Each column represents a different time point. From left to right, they are: 8 hours, 16 hours, and 24 hours post-infection. Only data from proteins in the intersection set (see Methods) are shown.

For PTM workflows, `ProteoMeter` requires protein abundance quantification and modified peptide intensities in tabular format. For LiP workflows, ProteoMeter requires tables containing LiP peptide intensities and trypsin-only peptide and protein quantification. In both workflows, `ProteoMeter` reads in a user-specified metadata table to interpret the columns of the protein and peptide quantification tables and group them according to control-treatment pairs, replicate groups, and batches. `ProteoMeter` additionally requires protein sequence information in fasta format. All data processing options can be specified in a configuration file in the human-readable, plain-text TOML format.

As an initial step, we provide simple visualization tools for assessing data quality. These include functions for computing principle component analysis (PCA) and constructing correlation plots. When running the data processing pipelines, users may specify which samples to omit as potential outliers. A configurable missingness is automatically applied to data during processing.

`ProteoMeter` conducts automatic median normalization of peptide and protein data, and will conduct row-mean centering batch correction if column batches are specified. It optionally applies protein abundance adjustments at the peptide level using either a traditional pooled-sample significance test or a direct matched-sample approach, depending on user input. In the former approach, proteins are tested for significant abundance changes between a control and treatment condition (the significance threshold is configurable, and can use either p-value or adjusted p-value). If a significant change is found, then the ratio of intensity means for the protein abundance measurements is used to scale the corresponding peptide groups. However, in this approach, variation in abundance quantification between replicates in the same group are retained at the LiP peptide level. Moreover, no abundance correction at all is performed when group-level abundance fold changes are not significantly different from random. This is more likely when there is naturally large variance in protein abundance, and it can lead to overestimating changes at the LiP protein level. Therefore, we implemented an optional alternative approach. In this latter approach, protein and peptide quantification samples are paired, and so each peptide sample is scaled according to the measured protein abundance in its partner sample. In this case, not only are the protein abundance variations between replicate samples in the same group removed, but also the changes of protein abundance between groups are removed to correctly quantify the site or peptide changes. This method cannot artificially remove measurement noise on average because such noise will not correlate between partners of a sample pair. This alternative abundance correction method is also supported by another independent work published recently [10].

Previous work [7, 11] has shown the utility of rolling up peptide level information to the single-residue level. For PTM experiments, aggregating the quantification of PTM to single site level significantly reduces the mismatches between different experiments and different runs [11]. It also increases the interoperability with the large number of published PTM studies conducted with antibodies that target a single PTM site. For LiP experiments, a recently published LiP data analysis tool, `FLiPPR` [7], applies a similar logic to the quantification of changes to “cut sites” to represent structural or exposure changes. Although `;FLiPPR` and `ProteoMeter` both aggregate the ion intensities of peptides sharing the same cut site, they differ in their aggregation approach. `FLiPPR` selects an aggregation function based on the pattern of missingness. In our implementation, we provide three different aggregation methods, namely *sum, mean*, and *median*, for directly aggregating the quantification regardless of the missingness. The default for both PTM and LiP peptide aggregation, *sum*, simply computes the sum of the (linear) intensity values; *mean* computes the mean of the log_2_ intensities; and the *median* computes the median of the log_2_ intensities.

In `ProteoMeter`, we perform residue-level roll-up on median-normalized peptide quantification before correcting for protein abundance and batch effects. After rolling up the quantification, we apply these secondary corrections directly at the residue level. We do this because protein abundance and batch effects do not necessarily have simple linear effects on peptide quantification. As such, it is typically impossible to conduct a perfect correction. Thus, the rolling up procedure can amplify any imperfections in the abundance and batch-effect corrections if done at the peptide level.

During and after normalization, roll-up, and correction, `ProteoMeter` automatically computes standard statistics– ANOVA and pairwise t-test p-values and adjusted p-values–at the protein, peptide, and site level (some of these are used during the correction procedures, while others are only used for data interpretation). The user may specify which comparisons to compute statistics for. We provide functions to visualize the significant difference between treatment conditions. These include functions for creating LiP barcode plots, volcano plots, and biplots. Data are output in a `pandas` data frame that can be easily saved in plain text tabular format.

### Application to MRC5 cells infected by HCoV-229E

In this section, we showcase the application of `ProteoMeter` to a study measuring responses of MRC5 cells to HCoV-229e infection at three time points: 8 hours, 16 hours, and 24 hours post-infection. We obtained nine measurement replicates for each time point and condition; four of these were used for multi-PTM analysis, and five were used for LiP analysis. In the multi-PTM analysis, we quantified CYS oxidation, LYS acetylation, and SER/THR phosphorylation. See Methods for further details of the experimental setup, data acquisition, and data analysis.

As detailed in Methods, we considered an intersection set composed of proteins for which we obtained measurements in both PTM and LiP experiments. In order to avoid undue sensitivity to missing values, we excluded proteins that did not contain at least one ProK digestion site positioned within an observed tryptic peptide. In total, our intersection set contains 928 proteins.

Comparing infected against control at each time point, and comparing each pair of time points for the infected condition, we observed 13,995 PTM sites (including oxidation, acetylation, and phosphorylation) and 419 ProK sites (1,331 tryptic peptides) significant at *q <* 0.05 (see Methods).

### Specific example

Among significantly post-translationally modified proteins with significantly altered ProK exposure, we selected Elongation Factor 2 (EF2) as a good candidate for further investigation and visualization (see Methods for selection criteria). We visualize the significantly altered residues in this protein in Figure 3. During GTP-dependent ribosomal translocation, peptidyl-tRNA and deacylated tRNA migrate between A, P, and E sites. EF2 is thought to catalyze the coordinated movement of these tRNAs, mRNA, and ribosome structure [12]. At the 16 hour post-infection time point, we observed a statistically significant increase in acetylation at several LYS residues coincident with changes in ProK digestion. Notably, the changes in digestion pattern in this example are not significant at the peptide level. However, averaging multiple peptides sharing a trypsin-only site increases statistical power and reveals statistically significant changes in digestion pattern. LYS acetylation has not been systematically studied in this protein, but Yao et al.[13] argued that LYS SUMOylation plays an important role in promoting proteolytic cleavage in EF2.

**Fig. 3.**
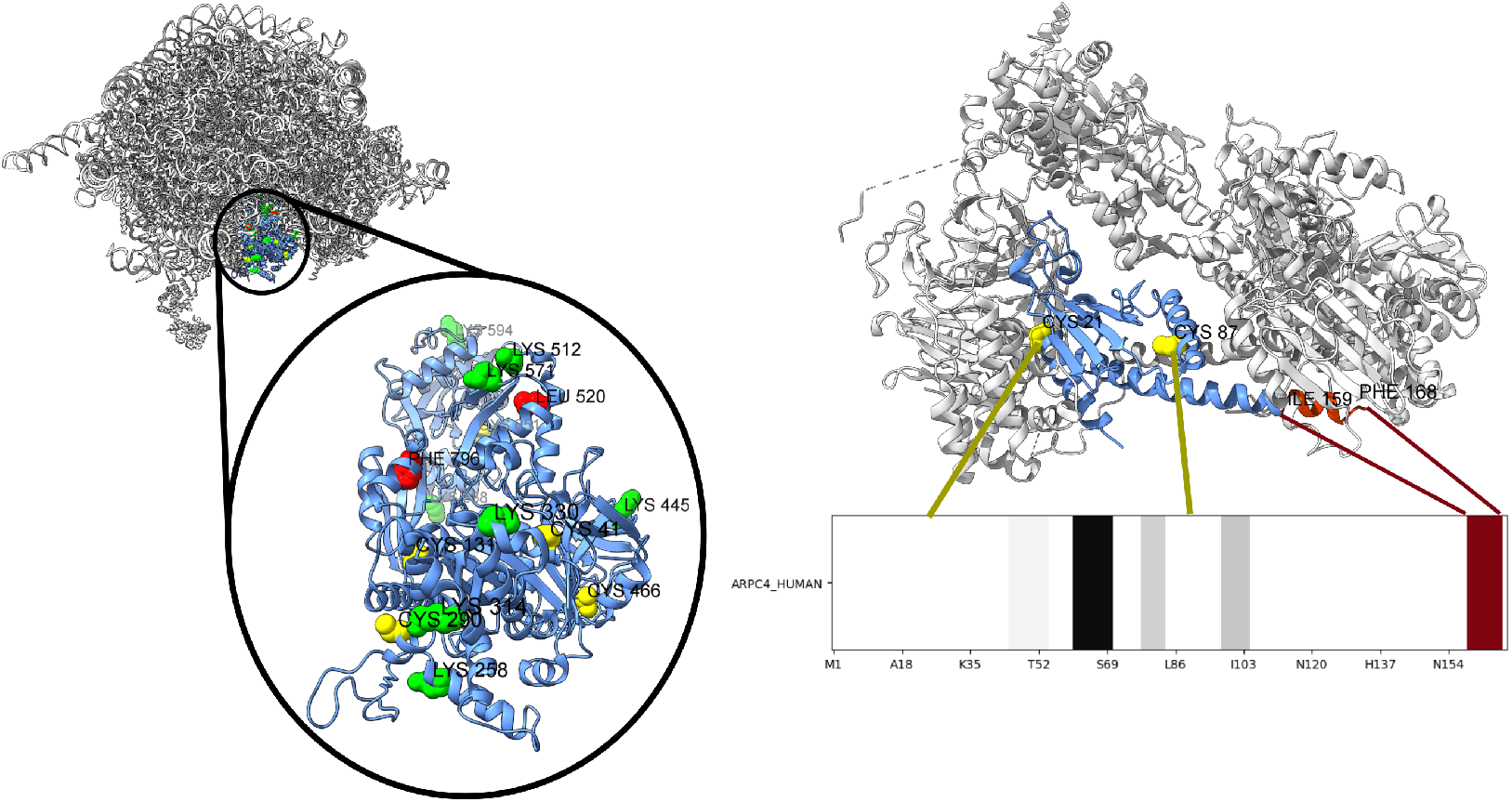
Example PTM and LiP results visualized on 3D structure. (**left**) PTM sites and ProK digestion sites on EF2 in the 80S ribosomal complex (PDB 4V6X [14]). The EF2 protein is highlighted in blue, with other proteins and RNA in white. Sites in red (LEU 520, PHE 796) indicate ProK digestion sites that differ significantly (*q <* 0.05) between the treated and control samples at the 16 hour post-infection time point. Similarly, green LYS residues indicate significant differential acetylation and yellow CYS residues indicate significant differential oxidation. Note that the ProK sites are not exposed in complex, consistent with the hypothesis that changes in the ProK site intensity are due to changes in binding. (**right**) Structure and barcode for ARPC4 in the Arp2/3 complex (PDB 6UHC [15]) at 16 hours post-infection (fold change relative to mock-infected control). ARPC4 is highlighted in blue, with significantly changed CYS oxidation sites (CYS 21 and CYS 87) shown in yellow. In the barcode plot, red indicates for significantly increased tryptic peptide abundance (decreased exposure) and gray for fold changes that are not statistically significant. In our barcode plot implementation in `Proteometer`, blue indicates significantly decreased tryptic peptide abundance (increased exposure), which did not occur in this protein at this time point. The darkness of the color indicates the magnitude of the fold change.

We can also examine putative structural changes by constructing a LiP barcode plot which shows measured tryptic peptides colored according to fold-change and significance. `ProteoMeter` implements functions for producing LiP barcode plots at the site or peptide level. As an example, we show in Figure 3 ARPC4 in the Arp2/3 complex with statistically significantly changed PTM sites and LiP tryptic peptide hilighted in color, alongside a barcode plot for the protein (all changes and significance calculated at 16 hours post-infection). ARPC4 is an acting-binding component of the Arp2/3 complex, which modulates cytoskeletal actin network formation and nuclear actin polymerization as part of DNA damage response mechanisms [16]. In this example, decreased exposure of the tyrptic peptide in the infected samples versus the mock-infected samples co-occurs with a decrease in oxidation of the two CYS oxidation sites (CYS 21 and CYS 87).

## Discussion

We have demonstrated `ProteoMeter`, a Python library designed to facilitate simultaneous, reproducible analysis of multi-PTM and LiP peptide and protein data with an emphasis on site-level quantification. We believe this emphasis will offer several benefits, especially when combined with peptide quantification. For instance, it aggregates multiple peptides for increased statistical confidence. In addition, Site-level information is more straightforwardly integrated into popular machine-learning paradigms such as graph neural networks. In the case of multi-PTM studies, it allows for statistical assessment of coordination or antagonism between individual protein modifications. In LiP analyses, it gives additional information about where structural changes have been observed to occur. In large, proteome-wide studies, we anticipate that comparing conclusions drawn from peptide-level and site-level statistics will be a fruitful way to further reduce false discovery, especially in LiP studies, which typically apply multiple testing corrections at the protein-level only [2, 17, 8, 10].

An advantage of `ProteoMeter` over other tools is that it integrates multi-PTM and LiP processing pipelines into a cohesive package, and uses human readable configuration file format for specifying options during data processing. This facilitates tracking data provenance during archival and iteration`ProteoMeter` exposes the full set of processing functions to the advanced user for incorporation into custom pipelines that deal with atypical experiment designs.

In this study, we have applied `ProteoMeter` to analyze the temporal, proteom-wide repsonse of MRC5 cells to HCoV-229e infection, highlighting differential PTMs and digestion pattern changes in EF2 and ARPC4. EF2 is plays an important role in regulation of ribosome function. Post-translational modification (SUMOylation) of LYS sites in EF2 have been shown be important for inhibiting its function [13], and our results highlight a possible complementary role for acetylation of these sites. ARPC4, through its actin-binding function in the Arp2/3 complex, modulates actin structure in the cytoplasm, which has previously been shown to undergo changes in lung fibroblast in response to infection [18]. Our results highlight a possible relationship between CYS oxidation in ARPC4 and formation of the Arp2/3 complex, though further investigation is required to definitively establish the nature of this relationship.

## Methods

### Experimental Overview

Here we provide a high-level summary of experimental procedures. Full details are available in the Supplemental Text. We obtained wild-type HCoV-229E from BEI resources for research purposes and used these to generate viral stocks. We infected immortalized human lung fibroblast cells (MRC5) cultured in alpha minimum essential medium. Cells were harvested at 8, 16, or 24 hours post-infection.

For PTM studies, five replicates were collected for each sample and processed following the procedure of [1]. We prepared 500*μ*L samples in deep-well plates and added an equal volume of absolute ethanol. Magnetic beads were conditioned and aliquoted into a bead plate. Samples were subjected to a digestion cocktail at 37^*°*^C for 3 hours. Cleaned peptide samples TMT-labeled and aliquoted for global, redox, phosphorylation, and acetylation enrichment prior to LC and IMAC analysis. All final peptide samples were analyzed on an ACQUITY UPLC (Waters) coupled with Q Extractive Plus mass spectrometer (Thermo Scientific).

For LiP studies, four replicates were collected for each sample and processed following the procedure of [2]. MRC5 cells in the LiP samples were lysed with three cycles of freeze-thawing and protein concentration was measured using BCA assay. Each sample was divided into a control part and a LiP part, each containing equal protein amounts. The LiP sample was treated with non-sepcific porease Proteinase K (ProK), while control parts of samples received an equivalent PBS volume, and all samples were incubated at 25^*°*^C for 1 minute followed by heat inactivation at 98^*°*^C for 5 minutes. Proteins were denatured by adding solid urea. Then, samples were reduced with DTT, alylated with IAA, and subjected to overnight digestion with LysC and trypsin. Samples were then acidified to 1% FA and desalted with C18 solid phase extraction. Final peptide concentrations were quantified using BCA assay and mixtures were analyzed using an Orbitrap Exploris mass spectrometer.

### Data analysis

Correlation plots generated in `ProteoMeter` revealed that one pair of infected-mock LiP samples at the 24 hour time point was poorly correlated with other replicates. We excluded this sample from our analyses. Additionally, for all peptide- and protein-level data, we excluded any rows (i.e. proteins or peptides) that did not appear in at least two replicates in the same group. We normalized all peptide- and protein-level data by centering the sample medians. Because protein abundance and peptide modification quantification measurements were conducted using paired samples, we applied this pairing to correct modification quantification for protein abundance, rather than aggregating within each treatment group as must be done in the absence abundance-modification sample pairing.

We conducted PTM measurements in two batches, and so we applied row-wise median-centering to correct for batch effects. We performed site-level roll-up using the sum of intensities prior to abundance and batch correction. We verified that batch effects were removed using PCA biplots generated automatically in `ProteoMeter` (see Supplemental Text).

After generating residue-level PTM and exposure data, we aggregated across measurements, discarding proteins that were not captured in all measurements. Among these remaining proteins, we identified those for which a significant change (*q <* 0.05) occurred both in PTM intensity and in ProK cleavage site exposure. Following established practice [2, 17, 10], we perform the protein-wise p-value adjustment of [2] for the LiP measurements to identify potential targets for follow-up investigation. To select an example for further investigation and visualization, we examined publicly available protein structures for proteins with fewer than 1,000 amino acids and which form complexes with experimentally measured structure. Of the proteins within this manually curated subset, we selected Elongation Factor 2 (EF2, UniProt ID P13639) as an illustrative example due to the completeness of the measured structure.

## Acknowledgments

The research described in this paper is part of the Predictive Phenomics Initiative at Pacific Northwest National Laboratory and conducted under the Laboratory Directed Research and Development Program. Pacific Northwest National Laboratory is a multiprogram national laboratory operated by Battelle for the U.S. Department of Energy under Contract No. DE-AC05-76RL01830.

