## supplementary file for "ProteoMeter: A pipeline for integrating multi-PTM and limited proteolysis data to reveal modification-structure coupling at the residue level"

### Supplemental Text

July 14, 2025

### 1 Methods

#### 1.1 Experiment details

*Viruses and Viral Titration.* Wild-type HCoV-229E was obtained from BEI resources for research purposes (NIAID, NIH: Human Coronavirus, 229E, NR-52726). Viral stocks were first generated (p1) in MRC5 (human lung fibroblast) and then p2 stocks were generated in Huh7 (liver epithelial) cells. Huh7 cells were also used to quantify viral titers following infection. Viral titers were quantified using standard plaque assay approaches [Sims et al., 2005]. Briefly, cells ($4.5 \times 10^5$ cells per well) were plated in 6-well dishes from Corning (Cat# 08-757-214). Viral samples were serially diluted, plated, and grown under 0.8% agarose and media on confluent monolayers. Plaques were visualized and counted following neutral red staining.

*Immortalized cell culture.* A549 (human lung epithelial cells, BEI resources: NIAID, NIH: Human Lung Carcinoma Cells (A549) Expressing Human Angiotensin-Converting Enzyme 2 (HA-FLAG), NR-53522) cells were cultured in high glucose Dulbecco's modified essential medium (DMEM; Gibco 11-995-073), containing 10% fetal bovine serum (FBS; Cytiva; SH3007003HI) and 1% Antibiotic-Antimycotic 100X (Anti-Anti; Gibco). MRC5 (human lung fibroblasts, ATCC CCL-171), were cultured in alpha minimum essential medium ($\alpha$-MEM; Gibco 12-561-056) containing 10% FBS and 1% Anti-Anti. HuH7 (human liver epithelial cells, Japanese Cancer Resource Bank JCRB0403) were cultured in low glucose DMEM (Gibco 11-885-092) containing 10% FBS and 1% Anti-Anti. Media serum concentrations were reduced to 4% prior to infection.

*Sample Processing for Proteomics.* A549 (PTM studies only) and MRC5 (PTM and LiP studies) cells were mock-infected or infected with HCoV-229E (MOI 3) and harvested at 8, 16, or 24 h post infection. Media from infected wells was collected at 8, 16, and 24 h post infection to determine HCoV-229E growth kinetics in both cell types. For LiP analysis, whole cell pellets were snap frozen and stored at $-80$ °C prior to being lysed using a hand-operated pellet-pestle homogenizer inside a biological safety cabinet. After measuring protein concentration through BCA, the lysates were subjected to LiP-MS. For PTM analysis, cells were washed once with $1\times$ phosphate buffered saline (PBS) and then washed again with PBS containing 100mM NEM for replicates 1 through

4. Replicate 5 for each condition (the total thiol sample) the final wash was performed with 1× PBS alone. Washes were removed and cells were frozen in their plates and stored at −80°C before harvest.

*Sample processing for multi-PTM proteomics.* Plates containing infected or mock MRC5 were lysed on ice with 200µL of 250 mM MES (pH 6.0) containing 5% SDS, 1% Triton X-100 with or without 100 mM NEM. Cells were collected and incubated in dark for 30min. DNA was sheered using a probe sonicator (Fisher Scientific Series 60 Sonic Dismembrator). Cell debris was removed by centrifugation at 13.000 g, 4 °C for 10 min. All samples were incubated at 55 °C, 850 rpm for 30 min. Automated sample cleanup and protein digestion was performed as described previously [Gluth et al., 2024, Leutert et al., 2019]. Briefly, the volumes of all samples in deep-well plates were adjusted to 500 µL, and 500 µL absolute ethanol was added (sample/binding plate). The magnetic beads were conditioned and aliquoted into a bead plate with a protein-to-bead ratio (w/w) of 1:5. Three wash plates contained 1 mL of 80% ethanol/well. The comb, bead, sample/binding and wash plates were placed onto KingFisher Flex (Thermo Scientific). The R2-P1 program was used to transfer beads to sample/binding plate, incubate for binding, and wash protein-bound beads for three times. The beads with proteins were transferred to a digestion plate containing 500 µL of digestion cocktail/well (1:100 trypsin and 1:100 Lys-C in 50 mL HEPES buffer, pH 7.7). Then digestion was conducted on a ThermoMixer at 37 °C, 850 rpm for 3h. The digested samples (the first elution) were collected on a MagnaBot FLEX 96 magnetic plate (Promega). 500 µL of 250 mL HEPES with 5% ACN (pH 7.7) per well was used to wash the beads and collected (the second elution buffer). Two elution was combined. Cleaned peptide samples were labeled with TMTpro 18-plex (Thermo Scientific) by incubation at room temperature, 850 rpm for 1h, followed by quenching with hydroxylamine. The TMT:peptide ratio (w/w) was 2.5:1. The labeled samples were cleaned up by C18 SPE desalting and aliquoted for global and redox, phosphorylation, and acetylation enrichment. Peptide level resin-assisted capture (RAC) of redox-modified peptides was conducted following the procedures published elsewhere using the thiol-affinity resin made in-house [Day et al., 2022, Guo et al., 2013, Gaffrey et al., 2021]. Global and redox peptide samples were fractionated on a nanoAcquity LC system (Waters) using a reversed-phase LC column (65 cm × 200 µm internal diameter packed with 3 µm Phenomenex Jupiter C18 particles) for 12 fractions [Li et al., 2021]. Automated IMAC phosphopeptide enrichment was conducted on an Agilent Bravo system as described by Abelin et al [Abelin et al., 2023]. The flowthrough from IMAC were collected and used for acetylation enrichment using PTMScan HS Acetyl-Lysine Motif [Ac-K] kit (Cell Signaling Technologies #46784) on KingFisher as previously described [Gluth et al., 2024]. All final peptide samples were analyzed on an ACQUITY UPLC (Waters) coupled with Q Exactive Plus mass spectrometer (Thermo Scientific) [Gluth et al., 2024].

*Limited proteolysis-based mass spectrometry (LiP-MS).* The MRC5 cell pellets were lysed with three cycles of freeze-thawing, and protein concentration in the lysates were measured using BCA assay. Each sample was divided into two parts- the control and the LiP sample- each containing equal protein amounts. The LiP sample was treated with non-specific protease, Proteinase K at an enzyme:substrate ratio 1:200. The control samples were received an equivalent volume of PBS. All the samples were incubated at 25 °C for 1 min, followed by heat inactivation at 98 °C for 5 min. The proteins were denatured by adding solid urea to achieve an 8 M solution. Subsequently, the samples were reduced with DTT (5 µM), alkylated with IAA (40 µM), and subjected to overnight digestion with LysC (enzyme:substrate = 1:100), and trypsin (enzyme:substrate = 1:100). Samples were acidified to 1% FA and then were desalted with C18 solid phase extraction. The final peptide concentrations were quantified using BCA assay, and peptide mixtures were analyzed using an Orbitrap Exploris mass spectrometer.

### 2 Quality Control

Figure 1 depicts the correlation between LiP samples at the peptide and protein level prior to further processing. Based on these plots, we removed the 24 hour post-infection mock-infected replicate number one and the 24 hour post-infection mock-infected replicate number one as outliers. We then recreated the correlation plots with these outliers placed in the last two rows and columns. Figure 2 depicts analogous plots for the PTM samples. No PTM samples were removed as outliers. However, we note pronounced batch effects in the correlation plots. We performed row-mean centering batch correction and compared PCA biplots before and after batch correction (Figure 3) to confirm that the dominant batch effect was successfully removed.

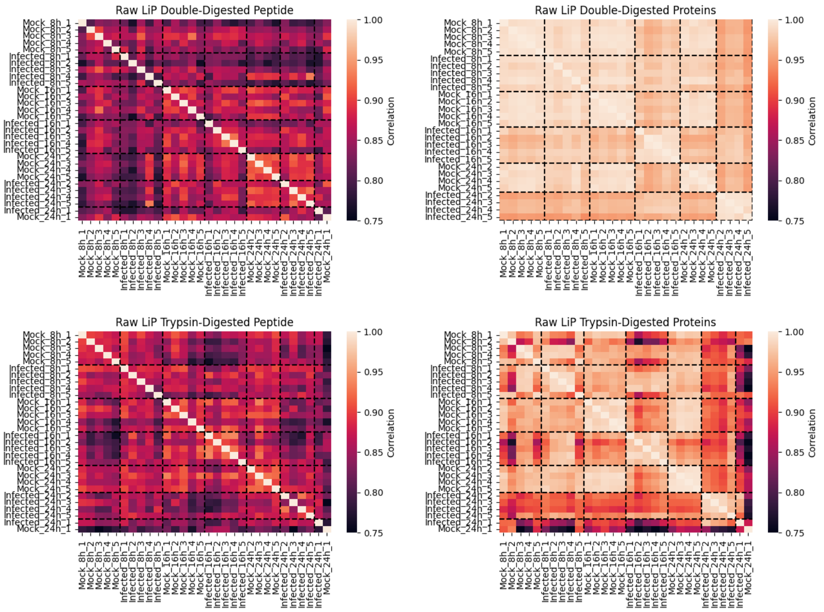


Figure 1: Pearson correlation plots for raw (input) LiP data at the protein and peptide level. Dotted lines separate replicate groups. The last two rows and columns are removed outliers.

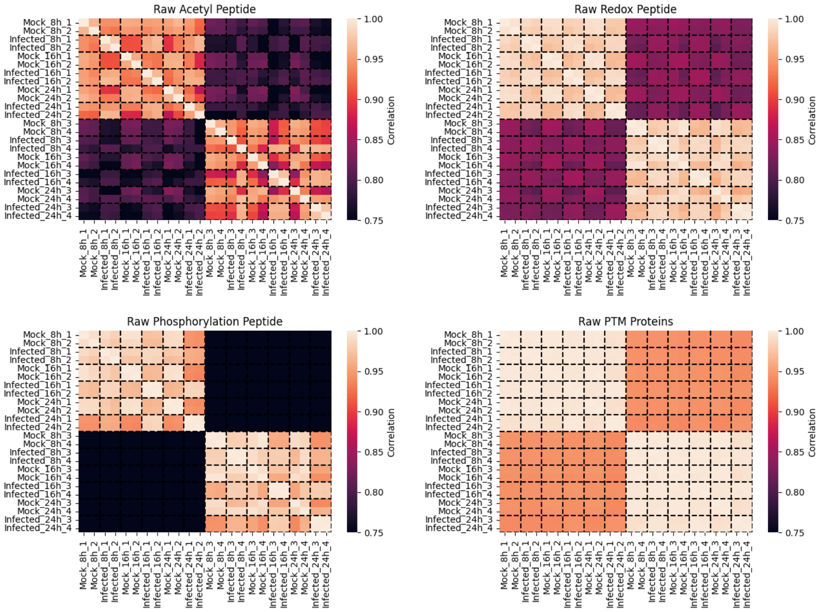


Figure 2: Pearson correlation plots for raw (input) PTM data at the protein and peptide level. Dotted lines separate same-batch replicate groups. Rows and columns are arranged so that the samples from the same batch are grouped together (there are two equally-sized batches). No samples were removed as outliers.

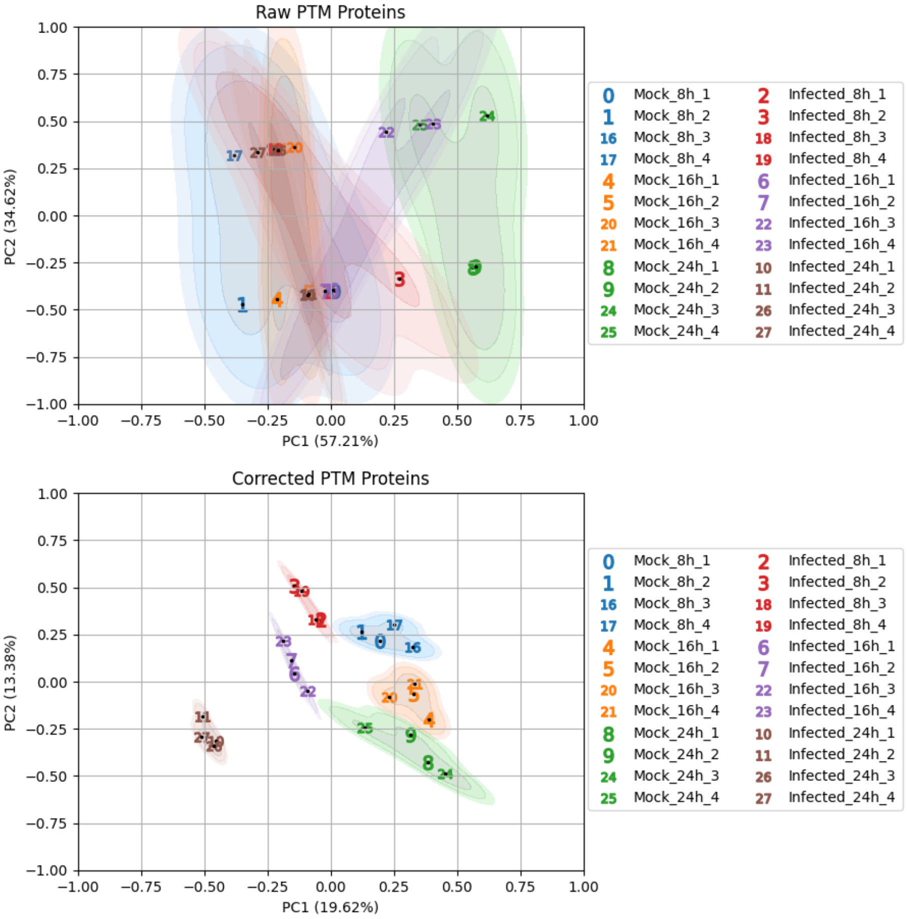


Figure 3: Comparison of PTM PCA biplots (protein-level) before (raw) and after (corrected) normalization and batch correction.

multiplexed quantification of protein thiol oxidation. *Journal of Visualized Experiments*, (172).

[Gluth et al., 2024] Gluth, A., Li, X., Gritsenko, M. A., Gaffrey, M. J., Kim, D. N., Lalli, P. M., Chu, R. K., Day, N. J., Sagendorf, T. J., Monroe, M. E., Feng, S., Liu, T., Yang, B., Qian, W.-J., and Zhang, T. (2024). Integrative multi-ptm proteomics reveals dynamic global, redox, phosphorylation, and acetylation regulation in cytokine-treated pancreatic beta cells. *Molecular Cellular Proteomics*, 23(12):100881.

[Guo et al., 2013] Guo, J., Gaffrey, M. J., Su, D., Liu, T., Camp, D. G., Smith, R. D., and Qian, W.-J. (2013). Resin-assisted enrichment of thiols as a general strategy for proteomic profiling of cysteine-based reversible modifications. *Nature Protocols*, 9(1):64–75.

[Leutert et al., 2019] Leutert, M., Rodríguez-Mias, R. A., Fukuda, N. K., and Villén, J. (2019). R2-p2 rapid-robotic phosphoproteomics enables multidimensional cell signaling studies. *Molecular Systems Biology*, 15(12):e9021.

[Li et al., 2021] Li, X., Day, N. J., Feng, S., Gaffrey, M. J., Lin, T.-D., Paurus, V. L., Monroe, M. E., Moore, R. J., Yang, B., Xian, M., and Qian, W.-J. (2021). Mass spectrometry-based direct detection of multiple types of protein thiol modifications in pancreatic beta cells under endoplasmic reticulum stress. *Redox Biology*, 46:102111.

[Sims et al., 2005] Sims, A. C., Baric, R. S., Yount, B., Burkett, S. E., Collins, P. L., and Pickles, R. J. (2005). Severe acute respiratory syndrome coronavirus infection of human ciliated airway epithelia: Role of ciliated cells in viral spread in the conducting airways of the lungs. *Journal of Virology*, 79(24):15511–15524.
